# PI3K activation in neural stem cells drives tumorigenesis which can be suppressed by targeting CREB

**DOI:** 10.1101/143388

**Authors:** Paul M. Daniel, Gulay Filiz, Daniel V. Brown, Michael Christie, Paul M. Waring, Yi Zhang, John M. Haynes, Colin Pouton, Dustin Flanagan, Elizabeth Vincan, Terrance G. Johns, Karen Montgomery, Wayne A. Phillips, Theo Mantamadiotis

**Affiliations:** Department of Pathology, School of Biomedical Sciences, University of Melbourne, Parkville, Australia; Stem Cell Biology Group, Drug Discovery Biology, Monash Institute of Pharmaceutical Sciences, Monash University, Parkville, VIC, Australia; Molecular Oncology Laboratory, University of Melbourne; Victorian Infectious Diseases Reference Laboratory, Doherty Institute; School of Biomedical Sciences, Curtin University, WA, Australia; Centre for Cancer Research, Hudson Institute of Medical Research, Clayton, Australia; Cancer Biology and Surgical Oncology Research Laboratory, Peter MacCallum Cancer Centre, Melbourne, Australia; Sir Peter MacCallum Department of Oncology, University of Melbourne, Parkville, 3010, Australia

**Keywords:** PI3K, PIK3CA, PTEN, CREB, WNT, WNT, brain cancer, glioma, GBM, mouse model

## Abstract

Hyperactivation of the PI3K signaling is common in human cancers, including gliomas, but the precise role of the pathway in glioma biology remains to be determined. Some limited understanding of PI3K signaling in brain cancer come from studies on neural stem/progenitor cells (NSPCs) where signals transmitted via the PI3K pathway cooperate with other intracellular pathways and downstream transcription factors to regulate NSPC proliferation. To investigate the role for the PI3K pathway in glioma initiation and development, we generated a mouse model targeting the inducible expression of a Pik3ca^H1047A^ oncogenic mutation and simultaneous deletion of the PI3K negative regulator, Pten, in NSPCs. We show that the expression of a Pik3ca^H1047A^ was sufficient to initiate tumorigenesis but that simultaneous loss of Pten, was required for the development of invasive, high-grade glioma. Mutant NSPCs exhibited enhanced neurosphere forming capacity which correlated with increased Wnt signaling. We also show that loss of CREB in *Pik3ca-PTEN* tumors led to a longer symptom-free survival in mice. Taken together, our findings present a novel mouse model for high-grade glioma with which we demonstrate that the PI3K pathway is important for initiation of tumorigenesis and that disruption of downstream CREB signaling attenuates tumor expansion.

## INTRODUCTION

Although our understanding of the cellular and molecular mechanisms controlling brain cancer pathogenesis has progressed over the last decades, the therapies available have not translated to improved patient outcome. This is exemplified in the most common and deadly high-grade glioma (HGG) in adults, glioblastoma (GBM), where post-therapy survival for most patients ranges between 8-14 months. Since the pioneering work by Holland and colleagues ^22^, genetically engineered mouse models have demonstrated that various gene combinations, overexpressing and/or targeting growth factors, growth factor receptors, oncogenic pathways and tumor suppressors in both brain progenitor or differentiated cells can initiate glioma development. These models, particularly those mimicking genetic alterations seen in patient tumors have proven to be invaluable in understanding the molecular events underlying the development of brain tumors, with the first models expressing constitutively activated mutant growth factor receptors (EGFR) or mutated key downstream signaling molecule (Ras, Akt) ^21, 22^. Numerous mouse models have since been generated using transgenic or gene modifying technologies targeting many genes ^1, 58^. Genes enabling the initiation of high-grade gliomas include combinations of *NF1, TP53, PTEN, AKT, Ras, INK4a/ARF* ^1, 2, 10, 43^. Many models have targeted glial cells, which are the cells enriched in human glioma ^10, 13, 21^. Other mouse models show that gene mutations in oligodendrocyte precursor cells or fully differentiated astrocytes or neurons can initiate glioma development ^16, 35^. The role of brain tumor stem cells (BTSCs) in fueling tumor growth has been demonstrated in a number of mouse models targeting genetic mutations/deletions in NSPCs ^1, 2, 22, 26, 35, 43, 49^ and highlighted by a study demonstrating that ablation of BTSCs inhibited tumor growth ^59^. With the better understanding of single gene or gene combinations driving HGG development, recent translational research has turned to modelling HGG by manipulating specific factors of cell signaling networks which can been targeted by pathway-specific drugs.

The signaling pathways driving HGG pathobiology overlap with those activated in many cancers and involve complex interactions between oncogenes and tumor suppressors. Most oncogenes and tumor suppressors are regulatory factors of the phosphoinositide 3-kinase (PI3K) and mitogen-activated protein kinase (MAPK) pathways. Both pathways activate a cascade of downstream kinases which lead to the activation of specific transcription factors, including the cAMP response element binding protein (CREB), which is upregulated in GBM and has a role in NSPC and GBM cell proliferation ^12, 45^. Emerging evidence suggest that oncogenic PI3K and MAPK signals converge with Wnt signaling to regulate cancer cell growth and proliferation^44^ and that in GBM, Wnt signaling has a role in cancer stem cell maintenance^48^.

When one considers the upstream components of the PI3K pathway including the epidermal growth factor receptor (EGFR) and the PI3K catalytic and regulatory subunits, PIK3CA and PIK3R1 as well as the pathway’s main negative regulator, phosphatase and tensin homolog (PTEN), up to 63% of HGGs, including GBM, exhibit an alteration in at least one of these genes ^31^. A recent study of cancer driver mutations shows that the catalytic subunit of PI3K, encoded by the *PIK3CA* gene, is amongst the three major oncogenic drivers in GBM; the other drivers are the *EGFR* and *TP53* ^52^. Indeed, *PIK3CA* mutations are reported to be present in up to 17% of all pediatric and adult brain cancer types with primary higher grade and treatment resistant brain tumors exhibiting the highest mutation rates ^31^. The full activation of the PI3K pathway not only requires enhanced catalytic activity to drive key downstream kinases such as Akt but also requires the inactivation of the lipid phosphatase activity of PTEN, which dephosphorylates phosphatidylinositol (3,4,5)-trisphosphate (PIP3) to the inactive PIP2 form^8, 14^. The precise role of PTEN in PI3K-driven HGG/GBM remains unclear, since recent discoveries demonstrate multiple PI3K-independent biochemical and cellular PTEN functions^6, 51^. Deletion of PTEN in neural stem and progenitor cells leads to enhanced proliferative activity in the stem cell niche but no tumor development ^19, 20^. However, *Pten* deletion in combination with *p53* and/or *Rb1* loss results in the development of astrocyte-derived high-grade tumors ^10^. An inducible viral expression system targeting NSPCs or glial cells, established the importance of platelet-derived growth factor-A (PGFA) in high-grade glioma initiation and the role of additional mutations of other genes, including *Pten* and *NF1*, in accelerating tumor development and determining the proneural GBM subtype^43^. The effects of pro-oncogenic mutations involving the PI3K catalytic (p110α) and regulatory (p85) subunits have not been investigated in mouse brain but combined activation of Ras and Akt in mouse neural progenitors resulted in glioblastoma formation ^22^, although neither oncogene alone led to tumor development.

In an effort to understand the key events in brain tumor development involving the PI3K pathway, we targeted the PI3K pathway in mouse NSPCs by conditional activation of an oncogenic mutation in *Pik3ca*, the gene encoding the PI3K catalytic p110α catalytic subunit of PI3K ^29^, in combination with the deletion of the PI3K pathway negative regulator, PTEN. To investigate the downstream transcriptional programs regulated by the PI3K pathway in our mouse model, we deleted CREB in *Pik3ca-Pten* mutant NSPCs and show that upon CREB loss tumor growth is attenuated via inhibition of mutant cell proliferation and subsequent conversion into a less aggressive tumor type.

## RESULTS

### Activation of *Pik3ca^H1047R^* expression in NSPCs is sufficient for tumor initiation but simultaneous deletion of *Pten* is necessary for the development of invasive tumors

The *Nestin-CreER^T2^* transgene used restricts Cre-recombinase expression to the sub-ventricular zone (SVZ) and the recombination only occurs following tamoxifen administration ^25^. First, mice heterozygous for a latent Cre recombinase (Cre)-inducible knock-in of the *Pik3ca*^H1047R^ mutation (*Pik3ca^H1047R-lox^*) ^29^ and/or two Cre-inducible *Pten* deletion alleles (*Pten^loxP/loxP^*) ^20^ were crossed with mice expressing a single (heterozygous) *Nestin-CreER^T2^* transgene. Further F1 crosses generated mice which were heterozygous for mutant *Pik3ca^H1047R^* (*Pik3ca^H1047R-lox^-Nestin-CreER^T2^*) and homozygous *Pten* deletion (*Pten^loxP/loxP^-Nestin-CreER^T2^*) (*Pten*^Δ^).

Control *Pik3ca^H1047R4ox^-Pten^loxP/loxP^* mice without the *Nestin-CreER^T2^* transgene but treated with tamoxifen, did not develop neurological symptoms, nor show evidence of tumor growth over the experimental time window of 200 days (Fig. 1A). Single mutant *Pten^Δ^* mice showed normal brain size with no overt abnormalities but did exhibit increased SVZ cellularity and proliferation compared to controls (Fig. 1A). Examination of single mutant heterozygous *Pik3ca^H1047R^* mouse brains between 50 to 150 days post-tamoxifen administration revealed the presence of tumors in the lateral ventricles (Fig. 1A).

**Figure 1.**
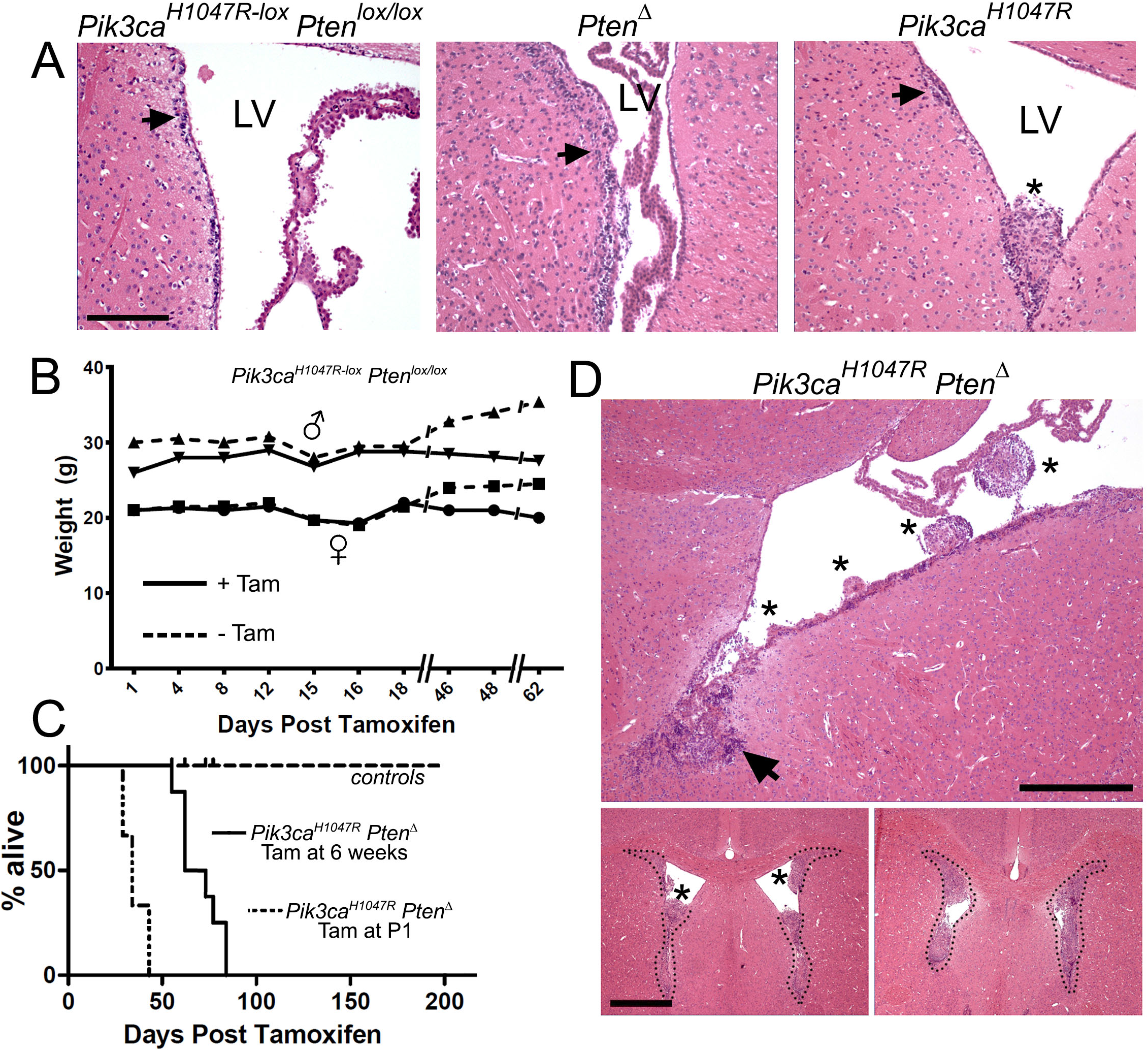
Combined *Pik3ca^H1047R^* oncogene activation and *Pten* deletion in neural stem/progenitor cells leads to the development of SVZ tumors. (**A**) H&E staining of brains from control mice (tamoxifen-treated *Pik3ca^H1047R^-Pten^lox/lox^* without *Nestin-CreER^T2^*) showing the SVZ layer (arrow) harboring NSPCs. Single mutant homozygous *Pten^Δ^*mice showed thickening of the SVZ layer (arrow). Single mutant heterozygous *Pik3cas^H1047R^* mice exhibited a normal SVZ (arrow) and a single tumor nodule (*) extending into the lateral ventricular space. Scale bar is 250μm. (**B**) Body weight of a representative cohort of adult mice (6 weeks of age) which received tamoxifen (Tam) at day 0. (**C**) Kaplan-Meier survival analysis shows that only tamoxifen-treated double mutant *Pik3ca***H1047R**-*Pten*^Δ^ adult (n=10) and newborn (P1) mouse (n=3) cohorts, compared to the control cohort (n=12). (**D**) H&E staining of a double mutant *Pik3ca*^H1047R^-*Pten*^Δ^brain shows the development of multiple tumors (*) along a single ventricular zone, each tumor at an apparent different stage of growth, which grow out from the neurogenic zone (arrows) into the lateral ventricles (LV) (Upper panel). Scale bar is 200 μgm. Anterior brain sections (lower panels) demonstrate almost complete occlusion of the ventricular space by the tumor tissue. Scale bar is 1mm.

Tamoxifen-treated double mutant *Pik3ca^H1047R-lox^-Pten^loxP/loxP^-Nestin-CreER^T2^* (hereafter referred to as *Pik3ca^H1047R^-Pten^Δ^*) 6-8-week-old mice, resulted in a completely penetrant (100%; 30 from 30 mice) neurological phenotype, apparent 55-90 days after tamoxifen administration. An early sign of ill health was a decline in body weight, compared to controls, in both male and female mice (Fig. 1B), followed by the appearance of progressively more severe neurological symptoms. The first neurological manifestation was ataxia, followed by sporadic seizures and a gradual increase in seizure frequency and length. Mice were culled when weight loss was more than 15% of the experiment start weight and/or when seizures occurred more than three times per day and at least one seizure lasted more than one minute. These criteria were deemed as the experimental endpoint. All tamoxifen-treated mice mutant mice reached the experimental endpoint between 55 and 90 days (Fig. 1C) and were subsequently culled. Furthermore, single mutant *Pik3ca^H1047R^* expression or *Pten* mice showed no body weight decrease, signs of brain dysfunction, nor reduced survival over 200 days (Fig 1B, C). When *Pik3ca^H1047R^-Pten^Δ^* mutations were activated in one day old newborn mice (P1) via tamoxifen treatment of the mother and transmission of the tamoxifen to pups via the mother’s milk, the pups began exhibiting severe neurological symptoms between 28 and 43 days (Fig. 1C), earlier than in adult mice treated with tamoxifen. P1 tamoxifen-treated mouse brains showed the presence of tumors, consistent with those seen in adult mice.

Histological examination of brains demonstrated that compared to control mouse brains, *Pik3ca^H1047R^-Pten^Δ^* mouse brains showed hypercellularity of the SVZ, with multiple tumor nodules protruding into the lateral ventricles (Fig. 1D, upper panel) and tumors completely filling the anterior lateral ventricles (Fig. 1D, lower panels). Further examination showed the presence of multiple hypercellular clusters lining the SVZ and tumor cell migration into the developing tumor nodules (Fig. 2A, B). Away from the lateral ventricles, tumor cells invaded the brain parenchyma and white matter tracts, including the corpus callosum (Fig. 2C). Apoptotic cells were scattered within tumors (not shown), lying amongst atypical, irregularly arranged tumor cells and blood vessels (Fig. 2D).

**Figure 2.**
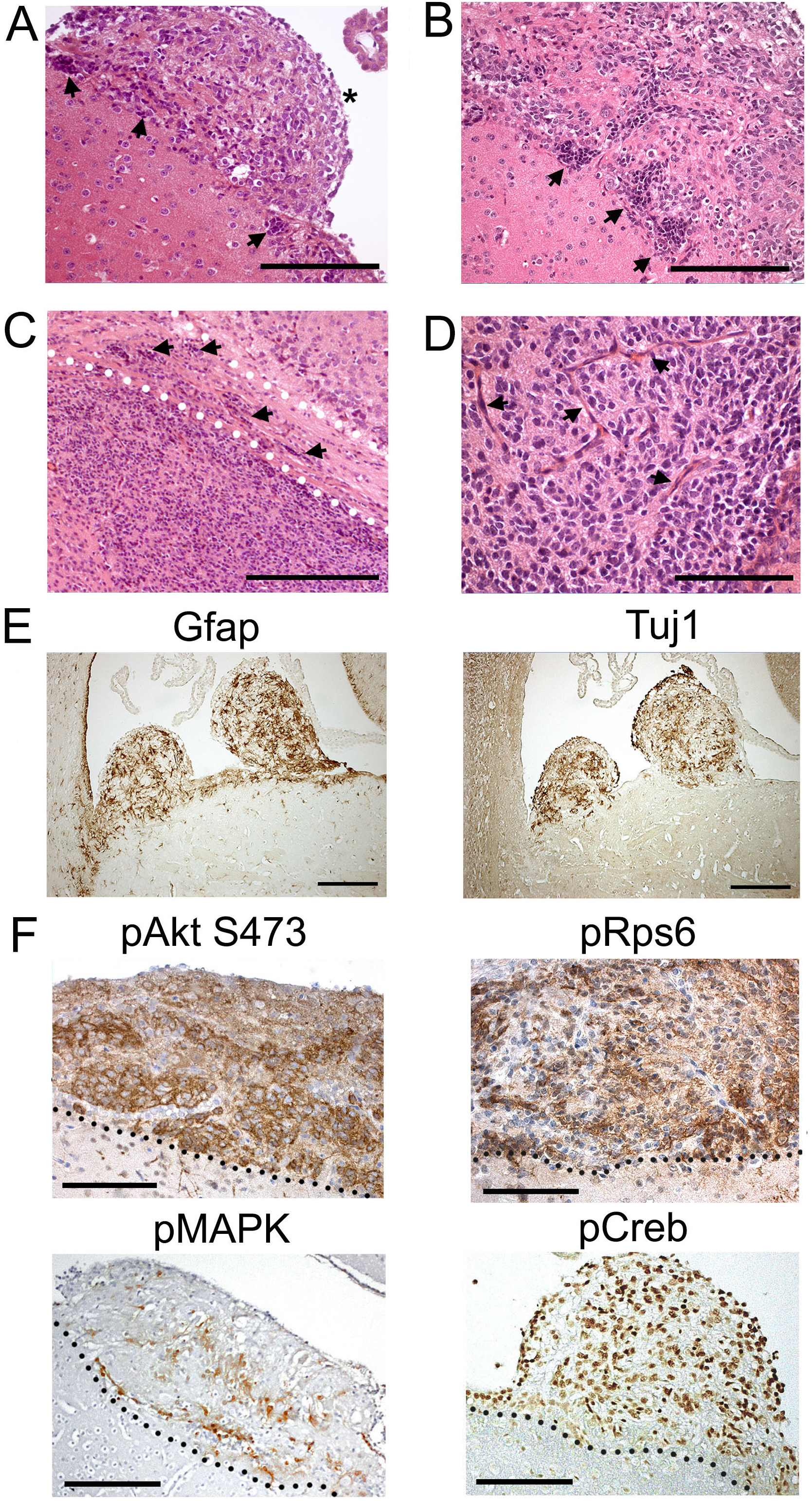
Mutant *Pik3ca^H1047R^-Pten^Δ^* NSPC initiated tumors exhibit features consistent with malignant astrocytoma. (**A**) A prominent tumor nodule (*) and hyperplastic germinal zones (arrows) (**A, B**). Tumor cell invasion (arrows) into the brain parenchyma and corpus callosum white matter tract (demarcated 30 by the dotted lines) (**C**); (**D**) tumor vascularization was observed in tumors. (**E**) Tumors expressed astrocytoma markers including GFAP and Tuj1. (**F**) Tumors exhibited elevated expression of pAKT(Ser473), pRpS6, pMAPK and pCREB. The dotted lines represent the SVZ-tumor interface, with tumors lying above the dotted line. Scale bars are: 200μm for (A), (E), (F); 100μm for (B); 500μm for (C); 20μm for (D).

Immunohistochemical analysis of *Pik3ca^H1047R^-Pten^Δ^* brains showed that tumors expressed glial fibrillary acidic protein (GFAP) and β-III-tubulin (Tuj1) (Fig. 2E) showing that the NSPCs from which the tumors developed could differentiate into glial-like and neuronal-like cells. pAKT (S473) and pRps6 expression were highly expressed in all tumor cells (Fig. 2F), evidence that the PI3K pathway was activated by the mutations. Activation of additional oncogenic signaling and transcriptional pathways was observed by examining phospho-ERK1/2 (pERK1/2) and phospho-CREB (pCREB) expression (Fig. 2F). Proliferating cell nuclear antigen (PCNA), Ki67 and nestin expression demonstrated that tumors also harbored proliferating, immature cells (Supplemental Fig.S1).

Overall, the histopathological analysis of the mouse brain tumors demonstrated heterogeneous features overlapping with both lower (II) and higher (III) grade astrocytic tumors. Specifically, the presence of atypical cells, evidence of cells undergoing apoptosis and tumor cell invasion into white matter tracts is consistent with low grade / WHO grade II astrocytoma, while the presence of blood vessels within tumors, suggests the presence of tumor phenotypes consistent with high-grade astrocytoma / WHO grade III.

### Mutant NSPCs exhibit enhanced proliferation and migration

Since mutant mice succumbed due to neurological complications associated with tumor growth, prior to broad dissemination of the disease, we assessed the invasive and functional characteristics of mutant cells in vitro. To obtain pure mutant NSPCs, SVZ tissue was used to isolate and propagate NSPCs from *Pik3ca^H1047R^-Pten^lox/lox^-UBC-Cre^ERT2^* mice ^30^ and Cre-mediated recombination was induced, in vitro, by the addition of 4-OH-T to the medium for 24h. Recombination in 4-OH-T treated cells was confirmed by PCR (data not shown) and loss of Pten protein (Supplemental Fig.S2), compared to untreated parental NSPCs and PTEN+ human GBM T98G cells. Mutant *Pik3ca^H1047R^-Pten^Δ^* neurospheres formed loosely aggregated rough-edged spheres (Fig. 3A) compared with parental vehicle treated control neurospheres. Moreover, mutant neurosphere cells showed the presence of more filopodia, compared to controls (Fig. 3A, B), a feature associated with a more invasive, malignant phenotype in GBM and some cancer cell lines ^41^. Incorporation of the thymidine analog, 5-ethynyl-2’-deoxyuridine (EdU) and flow cytometry analysis showed that *Pik3ca^H1047R^-Pten^Δ^* cells had a higher proliferation rate compared to parental Pik3ca-Pten wild-type cells (Fig. 3C). Mutant cells showed enhanced pAKT expression, in the presence or absence of EGF and bFGF (Fig. 3D). Correlating with enhanced proliferation, mutant cells exhibited increased expression of cyclins B1 and D1. Both control and mutant cells exhibited similar proportions of nestin-expressing cells (~97%) (Supplemental Fig. S2).

**Figure 3.**
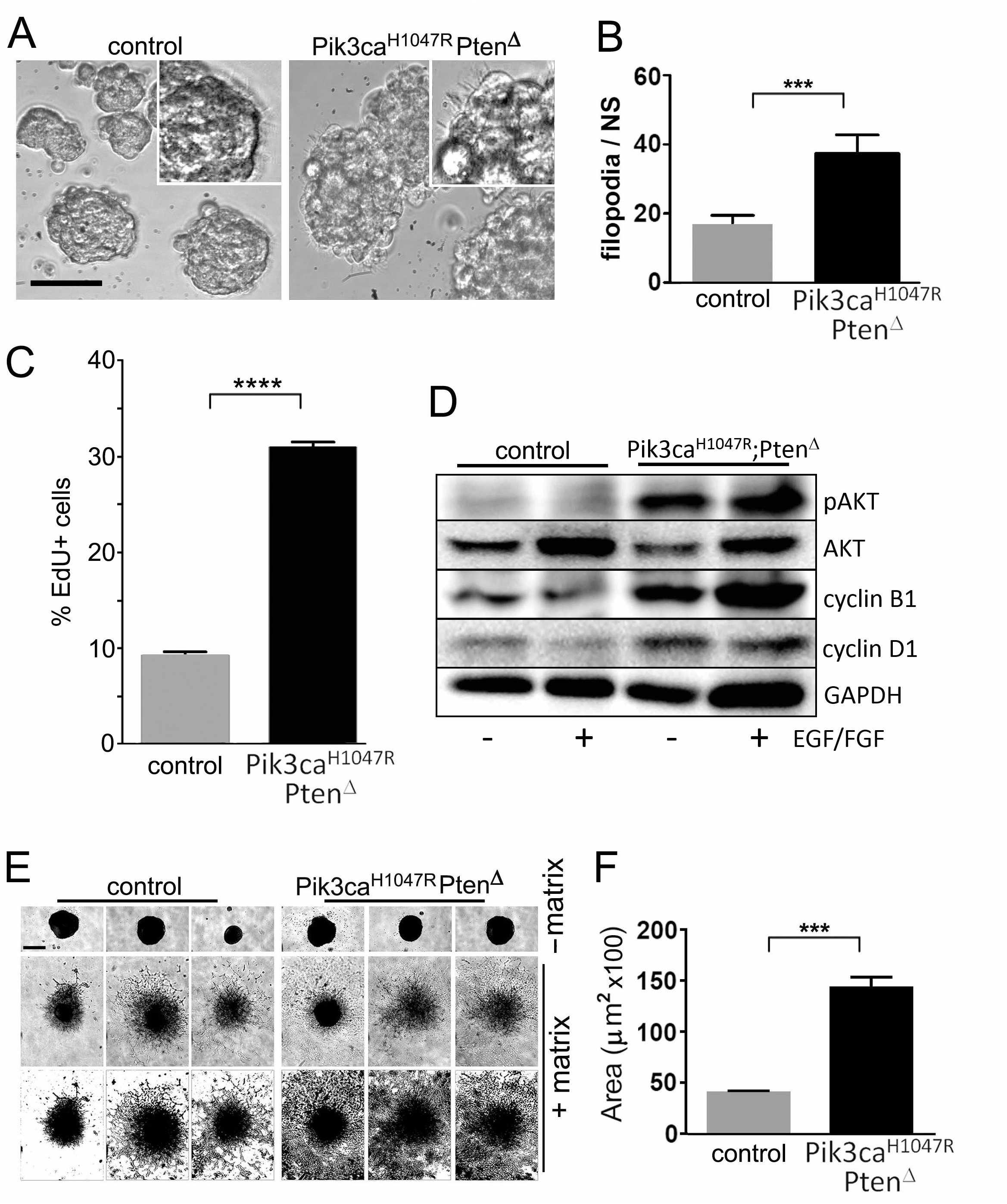
*Pik3ca^H1047R^-Pten^Δ^* mutations enhance the in vitro growth and migratory capacity of NSPCs. (**A**) Control and double mutant *Pik3ca^H1047R^-Pten^Δ^* NSPCs were grown in non-adherent neurosphere culture conditions. Control NSPCs form neurospheres with smooth circumscribed borders, while mutant NSPCs are larger forming mutant neurospheres exhibiting irregular edges. The inset, above right highlights part of a neurosphere with filopodia. Scale bar for the main images is 20μgm. (**B**) Mutant neurospheres (NS) showed the presence of more filopodia per cell, (mean ±SD, n=5; ***P<0.001, Student’s t-test). (**C**) *Pik3ca^H1047R^-Pten^Δ^* NSPCs exhibited enhanced proliferation, measured by EdU incorporation over 24h. ****P<0.00001. (**D**) Western blot analysis showing that *Pik3ca^H1047R^-Pten^Δ^* NSPCs express increased pAKT, cyclin D1 and cyclin D1, with or without EGF and bFGF-supplemented neurosphere medium. (**E**) Neurospheres seeded into a 96-well plate with or without invasion matrix for 48h show that in the presence of invasion matrix mutant NSPCs exhibit a higher invasive capacity compared to control NSPCs. The lower panels are “threshold” converted images (of the central (+ matrix) images) to enhance the contrast to show the extent of cell migration. Scale bar in top left panel of 31 (c) is 50μgm and applies to all images in the panel. (**F**) Quantitation of invasion expressed as area of spread (mean ±SD, n=5; ***P<0.001, Student’s t-test).

Seeding single neurospheres into a gel matrix showed that *Pik3ca^H1047R^-Pten^Δ^* NSPCs migrated to cover more than three times the area compared to control NSPCs at 48h (Fig. 3E,F), demonstrating that mutant cells have increased migratory capacity. Seeding cells at high density for 72h showed that mutant NSPCs detached and migrated away from spheres, unlike control NSPCs, which did not exhibit this behavior under the same seeding conditions (Supplemental Fig.S4).

### Mutant NSPCs exhibit enhanced sphere-forming capacity, persistent nestin expression and Wnt pathway activition

To determine the long-term proliferation differences between mutant and control NSPCs, cumulative growth was measured over 8 passages (8 weeks). *Pik3ca^H1047R^-Pten^Δ^* cells showed significantly higher cell number from week two onward and almost four times more cells by eight weeks (Fig. 4A). To elucidate the differences seen in cumulative growth, we next investigated the neurosphere-forming efficiency (a correlate of neural stem cell potential) differences between mutant and control NSPCs. Use of extreme-limiting dilution analysis (ELDA) ^24^ to measure neurosphere-forming efficiency, showed that in neurosphere culture conditions, the sphere-forming capacity of *Pik3ca^H1047R^-Pten^Δ^* cells (1 NSFU (neurosphere forming unit) / 1.55 cells seeded) was lower than control cells (1 NSFU / 2.01 cells seeded) (Fig. 4B). To test the inherent neurosphere forming stability of the NSPCs, we subjected the cells to alternating rounds of growth in neurosphere medium, serum-containing medium and a return to neurosphere medium to evaluate the ability of the cells in maintaining their neurosphere forming capacity under conditions which promote differentiation (Fig 4C). Flow cytometry analysis demonstrated that the NSPC marker, nestin, was still expressed in 70% of *Pik3ca^H1047R^;Pten^Δ^* differentiated cells compared with 52% of control cells after seven days (Fig. 4D). Returning differentiated cells to serum-free neurosphere conditions, revealed a significantly higher sphere-forming capacity in mutant cells (1 NSFU / 15.4 cells seeded), compared to control cells (1 NSFU / 36.0 cells seeded) (Fig 4E). Independent assessment by measuring nestin+, GFAP+ and Tuj-1+ cells grown on laminin for five days, supported the flow cytometric analysis with respect to an enhanced nestin+ cell stability under differentiating conditions. Mutant cells also exhibited higher GFAP+ cell numbers at day one but similar a number by day five, compared to parental wild-type NSPCs (Supplemental Fig. S5).

**Figure 4.**
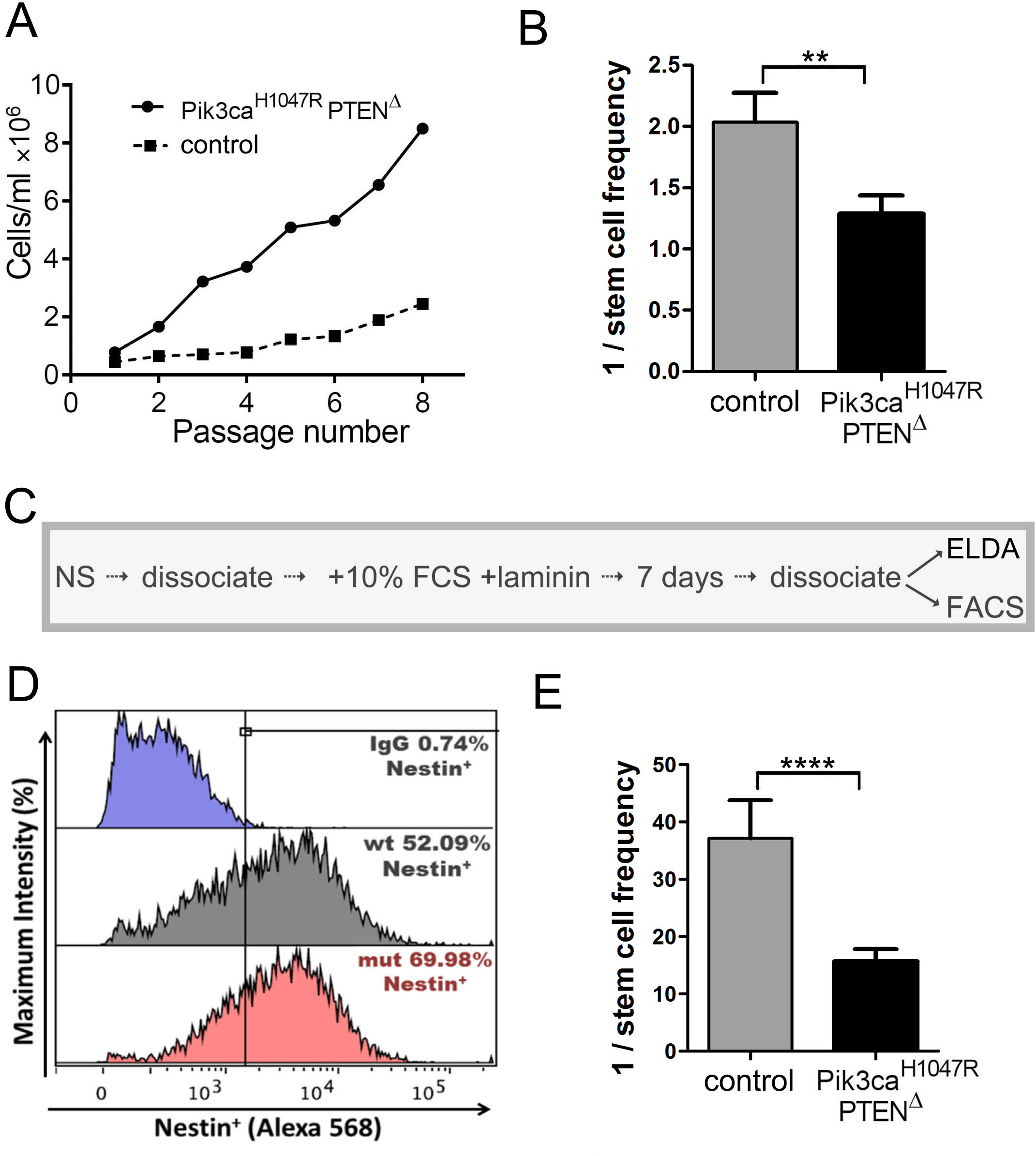
*Pik3ca^H1047R^-Pten^Δ^* mutant NSPCs exhibit enhanced sphere forming potential and maintenance of sphere forming potential under conditions promoting differentiation. (**A**) Cumulative cell number of control and *Pik3ca^H1047R^-Pten^Δ^* NSPCs over 8 passages (1,000 cells/ml seeded/1 passage/7 days). Cell counts were performed in triplicate from separate wells.(**B**) NSPCs seeded at low densities showed a selective advantage informing a neurosphere compared to control cells. n=3, Student’s t-test, **P<0.01. (**C**) The protocol flow used to test the maintenance of nestin expression and neurosphere forming capacity following differentiation.(D) Analysis of nestin expression by flow cytometry after differentiation for 7 days. After 7 days in differentiation conditions cells were returned to neurosphere medium and sphere-forming efficiency determined by extreme limiting dilution analysis (ELDA) (**E**), n=3, ****P<0.0001. See Materials and Methods for details.

Analysis of a panel of PI3K/AKT pathway-associated phosphoproteins in NSPCs grown in vitro, showed an upregulation of activation of numerous factors compared to parental control cells, including the factors: Akt, ribosomal protein S6 (rpS6), mTOR, GSK3α, GSK3β, p70S6 kinase, RSK1 and ERK1/2 (Fig. 5A). To further investigate the possible biological basis of the enhanced neurosphere forming capacity observed in *Pik3ca^H1047R^-Pten^Δ^* NSPCs, we postulated that Wnt signaling was upregulated in mutant cells, as cross-talk between PI3K and Wnt signaling is common in stem cells ^23^. Analysis of NSPCs showed that β-catenin was expressed at higher levels in mutant cell nuclei, compared to wild-type cells (Fig. 5B), indicating activation of Wnt signaling. Moreover, qRT-PCR demonstrated that the mRNA of Wnt pathway stem cell factors and targets was overexpressed in mutant NSPCs, compared to wild-type controls (Fig 5C). Further analysis demonstrated that neurosphere forming potential could be modulated by chemical inhibitors which modulate the Wnt pathway (Fig. 5E). CHIR99021 (‘CHIR’), a potent GSK3 inhibitor, and Wnt pathway activator, increased neurosphere forming potential of mutant NSPCs, where sphere per cells seeded capacity ratio, was 1:1 for CHIR-treated NSPCs, compared to 1:2.6 for vehicle (DMSO) treated cells. Upstream inhibition of the Wnt pathway, using IWP-2, which targets the membrane-bound protein, porcupine, and inhibits Wnt protein secretion, did not have a significant effect on sphere forming capacity, compared to vehicle treated mutant NSPCs, suggesting that the PI3K modulates Wnt signals downstream from porcupine and receptor activation.

**Figure 5.**
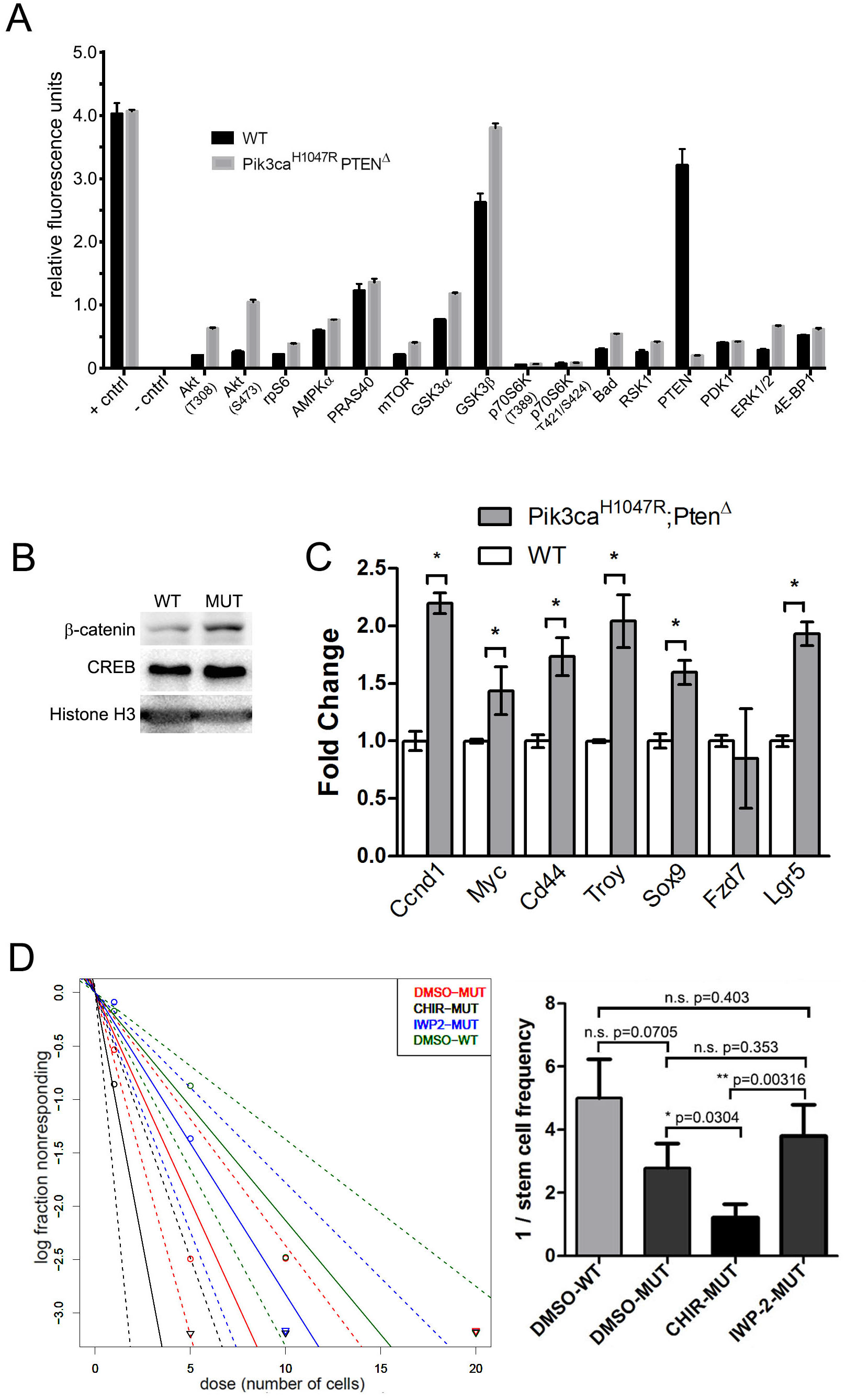
*Pik3ca^H1047R^-Pten^Δ^* mutant NSPCs exhibit enhanced PI3K downstream activity, MAPK activation, WNT pathway activation and WNT pathway-dependent sel-renewal capacity. (**A**) A PathScan^®^ Akt Signaling Antibody Array was used to measure the phosphorylation of a panel of PI3K/AKT-dependent factors in lysates of *Pik3ca^H1047R^-Pten^Δ^* mutant and control NSPCs. (**B**) Western blot showing increased nuclear β-catenin expression in *Pik3ca^H1047R^-Pten^Δ^* mutant NSPCs. Nuclear proteins, CREB and Histone-H3 were used as 32 loading controls (**C**) qRT-PCR analysis of a panel of WNT pathway factors showing upregulation of the WNT pathway in nuclei from *Pik3ca^H1047R^-Pten^Δ^* mutant cells. (**D**) Extreme limiting dilution analysis of NSPCs showing that WNT pathway activation by pharmacological inhibition of GSK3β using CHIR99021 increases neurosphere forming capacity of mutant NSPCs while WNT inhibition using IWP-2 reduces neurosphere forming capacity to wild-type levels. All error bars are S.E.M. from n=3. *p<0.05 or as indicated in.

### *Creb1* deletion suppresses malignancy of *Pik3ca^H1047R^-Pten^Δ^* tumors

Most cells in *Pik3ca^H1047R^-Pten^Δ^* tumors expressed pCREB (Fig. 2F) and this expression overlapped with nestin and GFAP expressing SVZ cells (Fig. 6A). We previously showed that mouse SVZ cells express activated pCREB ^15^ and human glioma tissue express high levels of both total CREB and pCREB ^12^. To determine the contribution of CREB in brain tumor development and growth, we used *CREB^lox/lox^* mice^39^ to generate triple mutant *Pik3ca^H1047R-lox^-Pten^lox/lox^-CREB^lox/lox^-Nestin-CreER^T2^* mice, resulting in deletion of CREB in addition to the *Pik3ca* and *Pten* mutations in NSPCs (*Pik3ca^H1047R^-Pten^Δ^-CREB^Δ^*, hereafter referred to as ‘triple mutant’).

**Figure 6.**
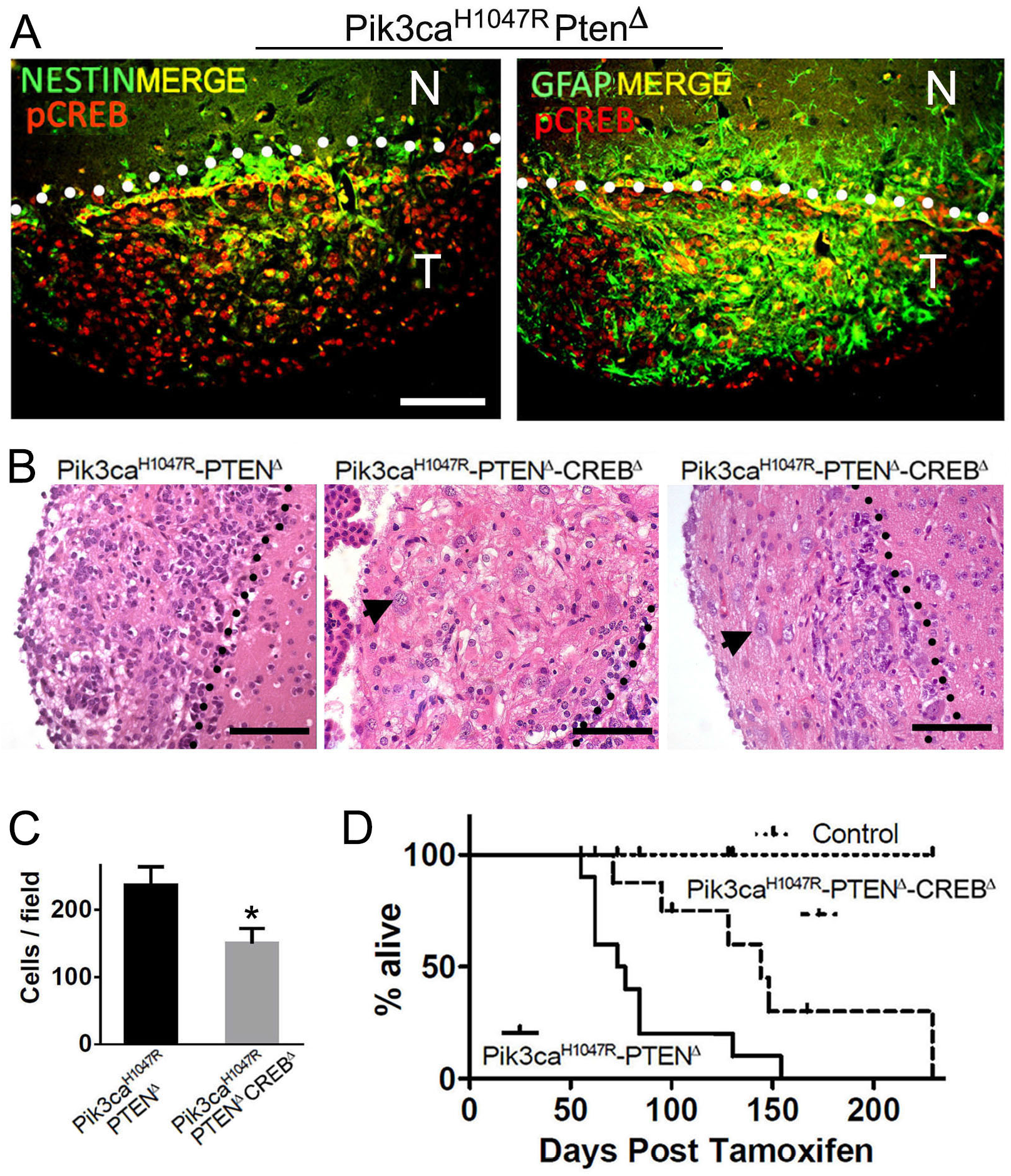
CREB deletion in *Pik3ca^H1047R^-Pten^Δ^* NSPCs increases survival by slowing tumor growth. (**A**) Immunofluorescence analysis of *Pik3ca^H1047R^-Pten^Δ^* tumors for expression of pCREB, the neural stem-cell marker, nestin, and glial-marker, GFAP. The SVZ-tumor boundary is demarcated by the dotted lines, with tumors lying below the line. Scale bars=100μgm. (**B**) H&E analysis showing differences in the cellularity of tumors derived from double mutant (*Pik3ca^H1047R^-Pten^Δ^*) (DM) and triple mutant (*Pik3ca^H1047R^-Pten^Δ^-CREB^Δ^*) mice. Triple mutant CREBΔtumors demonstrated the presence of large cells (arrow); absent in DM tumors. The SVZ non-tumor-tumor boundary is demarcated by the dotted lines; T=tumor, N=non-tumor. Scale bars = 100μgm. (**C**) Quantitative analysis of tumor cellularity between DM and triple mutant (DM+CREB^Δ^) (Mean ±SD, n=3; ** P<0.01, Student’s t-test). (**D**) Kaplan-Meier survival analysis of DM mice (n=10), compared to triple mutant *CREB^Δ^* mice (n=11) and control mice.

Triple mutant *CREB^Δ^* brain tumors exhibited the presence of large cells (Fig. 6B) and circumscribed growth with little or no invasion into non-tumor brain parenchyma (Supplemental Fig. 6A). We also observed a reduction in cell density in triple mutant *CREB^Δ^* tumors, compared to double mutant CREB wild-type tumors (Fig. 6B), which was confirmed by counting cells in DAPI stained sections (Fig. 6C), with triple mutant *CREB^Δ^* tumors exhibiting 65% cell density of *Pik3ca^H1047R^-Pten^Δ^* tumor cellularity. Notably, triple mutant *CREB^Δ^* mice were symptom-free for longer (median 125 days) following tamoxifen treatment, compared with double mutant *Pik3ca^H1047R^-Pten^Δ^ mice* (median 70 days) (Fig. 6D).

To explore the proliferative potential of *CREB^Δ^* NSPCs, in vitro proliferation assays were performed and showed that triple mutant *CREB^Δ^* NSPC proliferation was reduced compared with *Pik3ca^H1047R^-Pten^Δ^* (Supplemental Fig. 6B).

Consistent with the proliferation defect, cell cycle analysis showed that triple mutant *CREB^Δ^* NSPCs had fewer (5.9%) S-phase cells and more (80.2%) G0/G1 phase cells, compared to *Pik3ca^H1047R^-Pten^Δ^* (12.8% S-phase; 70% G0/G1-phase) NSPCs (Supplemental Fig. 6C). However, the sphere-forming capacity of *CREB^Δ^* NSPCs did not differ compared to *Pik3ca^H1047R^-Pten^Δ^* NSPCs (data not shown).

## DISCUSSION

The role of the PI3K pathway in driving malignant astrocytic brain tumors comes from murine models in which activation of retroviral vector-encoded AKT needs to be coexpressed with a second potent oncogene, such as Ras or BRAF, to trigger tumor development and growth ^22, 49^. To our knowledge, no studies have demonstrated a direct involvement of PI3K subunit mutations. Recent large-scale cancer sequencing data suggest that *PIK3CA* may be a key driver of malignant astrocytic brain cancer ^52^. Our study is the first to directly test the role of *Pik3ca* in brain tumor development and we demonstrate that expression of constitutive catalytically active Pik3ca^H1047R^ can initiate tumor growth when targeted to NSPCs, although simultaneous deletion of tumor suppressor Pten was necessary for the development of invasive malignant tumors, similar to that reported in other murine brain cancer models, where a second driver gene/tumor suppressor loss is necessary to trigger development of malignant tumors ^10, 26, 37^. In our model, homozygous loss of Pten only, did not lead to tumor development, although increased SVZ cellularity was observed, similar to the hyperproliferative neurogenic phenotype previously reported in *Pten^loxP/lox^-Nestin-Cre* mice ^3, 20^.

The NSPC tumors in the mice presented with highly variable, heterogeneous histopathological features, a characteristic of high-grade astrocytic tumors/GBM in young adult patients ^32^. The *Pik3ca^H1047R^-Pten^Δ^* tumors resemble some features commonly seen in patients with tuberous sclerosis, where mutations in the *TSC1* and *TSC2* genes, which encode factors downstream from the PI3K pathway, cause cortical layering defects, epilepsy, as well as the development of benign nodular subependymal tumors of the lateral ventricles ^11^. However, a *Tsc1* mutant mouse targeting NSPCS using a *nestin*-driven Tet-conditional transgene exhibits cortical disruption but no SVZ/ependymal tumors ^18^. Moreover, the data we present demonstrate the inherent invasiveness of the *Pik3ca^H1047R^-Pten^Δ^* tumor cells, where prominent tumor cell migration/invasion into the white matter occurred, an important feature of malignant astrocytomas and GBM. Furthermore, coexpression of both the glial marker GFAP and neuronal marker TuJ1 revealed in our model is significant, since this coexpression pattern in patient astrocytic tumors is associated with aggressive astrocytic tumors ^28^. Interestingly, tumors also exhibited rosette-like tumor cell arrangements tumors, which are features of both pediatric and young adult GBM ^32^.

One of the major roadblocks toward effective treatment of patients with high grade malignant brain cancers is the invasiveness of tumor cells into healthy brain tissue, which renders tumor removal by surgery almost impossible. Our data also shows that *Pik3ca^H1047R^-Pten^Δ^* NSPCs exhibit enhanced migratory properties and sphere-forming/stem cell-associated characteristics, consistent with previous studies demonstrating that PI3K pathway activation promotes GBM cell migration ^34^, neurosphere-formation and tumorigenicity of glioma stem cells (GSCs) ^54^.

Using extreme limiting dilution analysis, we found that *Pik3ca^H1047R^-Pten^Δ^* NSPCs were able to efficiently maintain nestin expression and sphere-forming capacity, even after forced differentiation. This implies that the *Pik3ca^H1047R^-Pten^Δ^* mutations stabilize stem cell properties, and, in a clinical setting, these mutations may impart stable, therapy tumor cell resistant characteristics and the capacity for tumor recurrence/relapse. Phospho-protein analysis of PI3K/AKT downstream targets revealed an array of upregulated factors. Notably, GSK3β, a key factor in the PI3K/AKT pathway, exhibited increased phosphorylation in mutant NSPCs, compared with wild-type control cells. GSK3P is also integral for activation of the Wnt pathway ^40^. Wnt and PI3K signaling are commonly co-activated in stem cells, which could explain the enhanced neurosphere forming capacity in *Pik3ca^H1047R^-Pten^Δ^* mutant NSPCs. Wnt signaling is a critical pathway for maintaining efficient self-renewal of NSPCs ^27, 55^ and is also implicated in maintenance of GSC self-renewal ^47, 57^. The clinical importance of Wnt signaling in patient gliomas is highlighted by the correlation of Wnt factors with independent markers of poor prognosis ^36^. Our data shows that *Pik3ca^H1047R^-Pten^Δ^* mutant NSPCs express high levels of all canonical Wnt pathway factors examined, except for the Wnt pathway receptor, Fzd7. Most of these genes examined have been reported to regulate NSPC and/or glioma cell biology. For example, the Leucine-rich repeat-containing G-protein-coupled receptor 5 (LGR5), plays a role in the maintenance and survival of patient-derived GSCs ^42^; Sox9 has a role in NSPC selfrenewal capacity ^50^; Myc inhibition disrupts mouse and human GSC self-renewal ^4^. Notably, cooperation between the PI3K and WNT pathways plays a key role in hematopoietic stem cell self-renewal ^46^, which may be similar to the cooperativity at play in NSPCs.

Cooperativity between the PI3K and MAPK pathways is suggested to be a requirement for GBM pathogenesis, explored in GBM in vitro and in vivo, using primary astrocytes with mutations in Kras and/or Pten ^53^. The authors observed that each individual mutation caused upregulation of both the PI3K and MAPK pathway and that inhibition of one of the pathways inhibited tumorigenic potential.

Although the oncogenic roles and mechanisms involving cytoplasmic signaling pathways, such as the PI3K and MAPK pathways are still being deciphered, the transcriptional programs activated by oncogenic mutations are less well understood. We and others have shown that the kinase-inducible transcription factor CREB and its activated form, pCREB are expressed in human astrocytic tumors, including the most deadly form, GBM ^5, 12^, and that CREB has a role in glioma cell proliferation ^12^. Analysis of *Pik3ca^H1047R^-Pten^Δ^* tumor brain tissue and NSPC lysates, showed that aside from the expected upregulation of the PI3K/AKT pathway factors, the MAPK pathway was also upregulated. Our previous work shows that a transcriptional convergence point of the PI3K and MAPK pathways is CREB ^12^. To explore the oncogenic role of CREB in vivo, we deleted CREB in *Pik3ca^H1047R^-Pten^Δ^* NSPCs. Strikingly, triple mutant *Pik3ca^H1047R^-Pten^Δ^-CREB^Δ^* mice were symptom-free for a longer period, compared with double mutant *Pik3ca^H1047R^-Pten^Δ^* mice. CREB deletion resulted in reduced tumor cellularity and a phenotypic shift of tumor cells toward a less malignant state, reminiscent of giant-cell GBM, which has a better prognosis compared with GBM ^33^. Moreover, no invasion into white matter tracts was observed in triple mutant *CREB^Δ^* brains. To our knowledge, this is the first in vivo data demonstrating that genetically targeted loss of CREB in tumor cells suppresses tumor growth and is consistent with studies correlating elevated CREB expression and activation with poor prognosis in many neoplasms including those in lung ^38^ and breast ^9^.

Overall, our data lead us to propose a model in which the PI3K pathway is critical and sufficient for reprograming normal NSPCs into tumor-initiating cells, in which the Wnt pathway is hyperactivated and contributes to NSPC self-renewal. As the resulting mutant tumor cells mature, the PI3K pathway activates a CREB-dependent transcriptome which contributes to tumor growth. MAPK signaling is also co-activated which probably contributes to CREB activation, to further fuel tumor cell proliferation. The putative efficacy of targeting CREB has led to the design of several experimental compounds which target CREB and can inhibit tumor cell proliferation in vitro and in vivo ^56^. Indeed, co-targeting archetypal oncogenic pathways, including PI3K, MAPK and WNT, as well as transcription factors such as CREB, may prove to be effective therapeutic approaches for difficult to treat cancers, including HGGs, since transcription factors are less prone to direct mutations, thus circumventing the development of drug resistance ^17^.

## Materials and Methods

### Ethics statement

Experiments in mice were carried out with the approval of The University of Melbourne, School of Biomedical Sciences (AEC No 1112336.1) and Peter MacCallum Cancer Center (AEEC No. E406) Animal Ethics Committees.

### Mouse maintenance and husbandry

Mice were housed at the Biomedical Sciences University of Melbourne animal house in a pathogen-free environment with food and water freely accessible. Environmental supplementation of the cages was limited to necessary bedding materials to standardize conditions between cages and mice. *Pten^lox/lox^* (c;129S4-Ptentm1Hwu/J) were from The Jackson Laboratory. *Pik3ca^H1047R-lox^* ^29^, *CREB^lox/lox^* ^39^, and *Nestin-CreER^T2^* ^25^ mice were generated, as previously described. All mice were on a C57BL/6 background.

### In vivo tumor induction

Control (*Pik3ca^H1047R-lox^-PTEN^lox/lox^*), double mutant (*Pik3ca^H1047R-lox^-PTEN^lox/lox^-Nestin-CreER^T2^*) or triple mutant (*Pik3ca^H1047R-lox^-PTEN^lox/lox^-CREB^lox/lox^-Nestin-CreER^T2^*) mice were treated with tamoxifen at 6-8 weeks of age, as described previously ^7^.

### NSPC isolation and culture

Adult (6-10 week) control (*Pik3ca^H1047R-lox^-PTEN^lox/lox^*), double mutant (*Pik3ca^H1047R-lox^-PTEN^lox/lox^-Nestin-CreER^T2^*) or triple mutant (*Pik3ca^H1047R-lox^-PTEN^lox/lox^-CREB^lox/lox^-Nestin-CreER^T2^*) mice were culled immediately before harvesting. Brains were placed in ice cold PBS before dissection of the SVZs and NSPCs were isolated as previously described ^15^. NSPCs were grown in Dulbecco’s modified eagle medium (DMEM) (Invitrogen, USA) with the addition of 2% B27 supplement (Invitrogen), 20 ng/mL basic fibroblast growth factor (b-FGF, Sigma, USA), and 40 ng/mL epidermal growth factor (EGF, Sigma). NSPC maintenance and passaging was performed as previously described ^15^.

### In vitro controls

For experiments involving NSPCs, mutant NSPCs were generated by transient (24h) treatment with 0.02mg/ml 4-OH-T (in methanol) to induce Cre-mediated recombination of floxed alleles, while control cells were derived from the same genotype but treated with vehicle only (methanol). 4-OH-T was removed after 24h and never included in the medium in further experiments / assays.

### Cell cycle analysis

Cells were dissociated and then resuspended in PBS. Cells were then fixed by adding cell solution dropwise into ice cold 100% ethanol. Fixed cells were kept at -20°C until day of analysis where they were washed 3x with PBS and then incubated with DAPI staining solution (5μg/mL; Thermo Fisher Scientific, MA USA) for 15 minutes on ice. Following incubation, cells were washed, resuspended in PBS and immediately run on a LSR Fortessa using the UV laser (405nm). Cells were first gated with forward scatter width (FSW) vs DAPI (405nm) to exclude doublets and debris. A histogram of DAPI vs count was generated and analyzed on the flow cytometry analysis software FlowLogic (version 6) to identify the population (%) of cells in each of the cell cycle phases.

### Proliferation assays

Cells were plated onto a 96 well plate and grown under required treatment conditions. On the day of analysis, Resazurin solution (Sigma, MO, USA) was diluted in appropriate media and added to wells to obtain a 10% v/v solution before incubation for at least 3 hours at 37°C. Plates were analyzed using the EnSpire Plate Reader (PerkinElmer).

### Migration assay

Single neurospheres grown in DMEM-F12 with growth factors and supplements (see above) for 48h were placed into a 96-well plate (1 sphere/well) and assayed using the Trevigen 96-well 3D spheroid BME cell invasion assay kit (Bio Scientific, Australia), according to the manufacturer’s protocol. Briefly, neurospheres were incubated either with (test) or without (control) addition of invasion matrix for 48h. Spheres were photographed on an inverted microscope and invasion area was calculated by measuring the area of invasion by subtracting the control neurosphere area using Image J (http://rsb.info.nih.gov/ij/index.html).

### Extreme Limiting Dilution Analysis (ELDA)

Neurospheres were dissociated and plated in suspension media at decreasing cellular densities (20, 10, 5, 1 cell) using 24 wells per cell density condition. Wells were imaged 7 days after plating and the number of wells with one or more sphere, greater than ~20μm diameter, were scored. Data was analyzed as previously reported ^24^ using the online ELDA software hosted by WEHI (http://bioinf.wehi.edu.au/software/elda/). Cells were plated at 200,000 cells/9.61cm^2^ in DMEM-F12 with 10%FCS and cultured for 7 days before subsequent analysis.

### Flow cytometry analysis

Cells were harvested to single cell suspension as described above and resuspended in PBS. Cells were then fixed by adding cell solution dropwise into ice cold 100% ethanol. Fixed cells were kept at -20°C until day of analysis when they were washed 3x with PBS and then incubated for 30 minutes in blocking solution (PBS supplemented with 2% (v/v) normal goat serum and 0.1% (v/v) Tween-20). Cells were spun down and supernatant discarded before nestin antibody (Millipore) diluted in blocking solution (1:200) was added to cells. The primary antibody was left on for 60 minutes before a wash in PBS and staining cells with an anti-mouse Alexa-Fluor 568 secondary antibody (1:500) (ThermoFisher) for 10 minutes. DAPI staining solution was added to cells after the 10-minute incubation had elapsed and left to incubate for a further 5 minutes on ice. Following a final wash in PBS, cells were resuspended in 100μl of PBS and immediately run on a LSR Fortessa. Cells were gated with forward scatter width (FSW) vs DAPI (405nm) to exclude doublets and debris. Flow data was analyzed using the analysis software FlowLogic (version 6) to identify the population (%) of cells in each of the cell cycle phases.

#### Statistical analysis

Data were analyzed using the Student’s t-test and were presented as mean ± standard deviation (SD) or standard error of the mean (SEM), as indicated in relevant data/sections. P<0.05 (*) was considered as significant.

## CONFLICT OF INTEREST

The authors declare no conflict of interest.

## ACKNOWLEDGMENTS

We thank Rob Ramsay, Frederic Hollande, Andrew Allen, Daniel Gough, Jason Cain, Ryan Hutchinson for helpful scientific discussions. We thank Tina Isaakidis for proofreading. Special thanks to Sarah Louise Taverner for dedicated animal care and monitoring throughout this project, as well as to Teresa Drever, Jessica Sturrock, Michelle Williams and Marica Kesar for running and managing the excellent animal facility. Daniel Blashki and Vanta Jameson for flow cytometry assistance. We thank the Department of Pathology and The CASS Foundation Science & Medicine Grant #6236 for financial support. WAP was supported, in part, by project grant #1080491 from the National Health and Medical Research Council (NHMRC) of Australia. We would also like to acknowledge the donors of the tumor-derived sample data within the TCGA, Rembrandt GBM Datasets.

Supplementary Information accompanies the paper on the *Oncogene* website (http://www.nature.com/onc)

